# A chromosome-level reference-quality genome of *Punica granatum* L

**DOI:** 10.1101/2024.06.02.596999

**Authors:** Ming Yan

## Abstract

Pomegranate (*Punica granatum* L.) is one of the most ancient edible fruit tree species. Here we reported a new chromosome-level genome assembly and annotation of sour pomegranate. We assembled the genome with a size of 331.47 Mb and used BUSCO to estimate the completeness of the assembly as 98.8%. More than 97.40% of sequences in the final assembly were anchored to 8 pseudochromosomes, higher than the corresponding percentages for the existing reference genomes ‘Tunisia’ (92.62%). Using a combination of de novo prediction, protein homology and RNA-seq annotation, 29,326 protein-coding genes were predicted. We re-annotated the protein-coding genes of five other published pomegranate genomes using the same annotation method. We constructed the pan-genome of pomegranate using protein-coding genes, integrating data from our newly assembled genome and five other published genomes. The pan-genome was composed of 28,314 gene families, of which 68.96% were core genes, 30.00% were dispensable genes, and 1.04% were private genes. The chromosome-level reference genome of sour pomegranate would be valuable resource for research and molecular breeding of pomegranate.

## Background & Summary

Pomegranate (*Punica granatum* L.) is one of the most ancient edible fruit tree species native to central Asia ^1^. Popularity of pomegranate has increased tremendously over the past decade because of its functional and nutraceutical properties ^2^. Its fruits are rich in the phenolic compounds, including punicalagin, gallic acid, and ellagic acid, which are potentially beneficial in preventing cardiovascular disease and diabetes ^3^. In addition to being eaten fresh, pomegranate fruits are used in food industries to prepare juices, wines, and jams ^4^. The oil obtained from pomegranate seeds is known for its functional and beneficial health properties ^5^. Apart from its commercial importance, pomegranate has become an attractive system for studying several valuable biological features, such as color formation in the fruit peel and aril ^6^, phenotypic variations between soft- and hard-seeded varieties ^7,8^, and the mechanism underlying ovule sterility ^9-11^. Development of high-quality reference genome would be very useful for understanding the molecular mechanisms underlying these valuable biological features. The first draft whole genome sequence for ‘Dabenzi’ pomegranate was published in 2017 ^12^. The total length of this assembly was 328Mb, with 29,229 annotated gene models. This assembly indicates that a Myrtales lineage-specific whole-genome duplication event occurred in the common ancestor before the divergence of pomegranate and eucalyptus. Subsequently, ‘Taishanhong’ pomegranate genome ^13^ was released, which resolved the previously debated taxonomic status of the genus Punica, and reclassified it in the family Lythraceae. In 2019, a high-quality chromosome-scale genome of the soft-seeded pomegranate cultivar ‘Tunisia’ was published ^14^, which provided an attractive candidate region for further investigation for the genetic factors contributing to the divergence between hard- and soft-seeded pomegranate cultivars. Recently, two draft genome assemblies of the main Indian cultivar ‘Bhagwa’ were reported ^15,16^. The increasing availability of genome sequences of different pomegranate cultivars has facilitated advances in basic research, comparative, and evolutionary genomics studies of *P. granatum* ^17^. Here we reported a new chromosome-level genome assembly and annotation of sour pomegranate landraces from Xinjiang, China (hereafter called yechensuan) using Oxford Nanopore. A total of 27.9 Gb of Nanopore sequences with N50 read length of 51.06kb was generated, covering approximately 78x of the yechensuan genome with an estimated size of 355.86 Mb. The Nanopore long reads were initially assembled into 66 contigs with an N50 of 16.78Mb (Table 1). After excluding putative chloroplast and mitochondrial genome assemblies, the contigs were then scaffolded using RagTag based on *Punica granatum* cv ‘Tunisia’ reference genome. A final total of 66 contigs were arrayed into 24 scaffolds with an N50 of 41.86Mb, which are significantly better than the corresponding values of previously published pomegranate genome assemblies. The final assembly resulted in 8 pseudomolecules covering ∼ 97.40% of the genome, having a final assembled genome size of 331.47M (Fig 1A). Busco was used to evaluate genomic completes. 98.8% of the core conserved plant genes were found complete in the genome assembly. The LAI value was 12.30, which indicated that out newly assembly reached the ‘reference’ level (LAI > 10). Further evaluation using Merqury revealed a consensus quality score (QV) of 32 for the yechensuan assembly. Collectively, these results indicated that the yechensuan genome assembly is of high quality with substantially improved contiguity and completeness compared to the five previously reported pomegranate assemblies. Based on the species-specific TE database constructed by RepeatModeler2, the repeats were annotated using RepeatMasker. Finally, 50.26% of our newly assembled genome was masked as repetitive sequence. Long terminal repeat retrotransposons (LTR-RTs) representing the most abundant class of TE (Fig. 1C). Using a combination of de novo prediction, protein homology and RNA-seq annotation, 29,326 protein-coding genes were predicted, with an average coding sequence length of 3,050 bp and an average exon length of 219 bp. A total of 97.7% of completely assembled genes was assessed by the BUSCO analysis.

**Table 1.** Statistics of sour pomegranate genome assembly and annotation.

|  |  |
| --- | --- |
| Total size of assembled scaffolds (MB) | 331.47 |
| Number of scaffolds | 24 |
| N50 scaffold length (Mb) | 41.87 |
| Longest scaffold (Mb) | 68.22 |
| Number of contigs | 66 |
| N50 contig length (Mb) | 16.78 |
| Largest contig (Mb) | 21.89 |
| GC content (%) | 41.08 |
| Number of gene models | 29,326 |
| Mean coding sequence length (bp) | 3,050 |
| Mean number of exons per gene | 5.4 |
| Mean exon length (bp) | 219.2 |
| Complete BUSCOs (Genome) | C:98.8%[S:96.6%,D:2.2%],F:0.2%,M:1.0%,n:1614 |
| Complete BUSCOs (Predicted genes) | C:97.4%[S:95.8%,D:1.6%],F:1.1%,M:1.5%,n:1614 |
| Total base of Illumina reads (Gb) | 14.2 |
| Total base of Nanopore reads (Gb) | 27.9 |

\* Complete BUSCOs (C); Complete and single-copy BUSCOs (S); Complete and duplicated BUSCOs (D); Fragmented BUSCOs (F); Missing BUSCOs (M)

**Figure 1.**
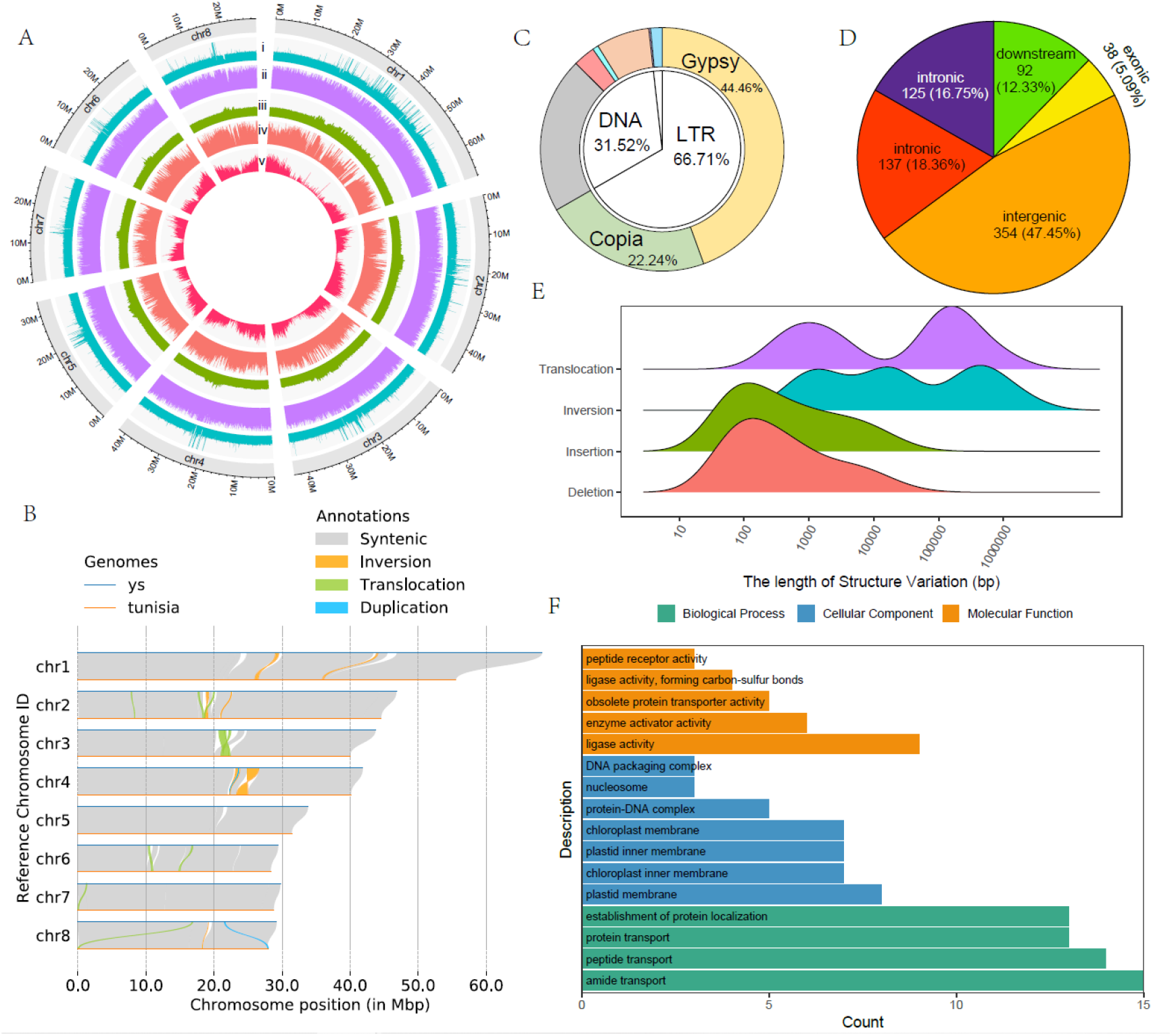
Genomic landscape of Punica granatum cv. ‘yechensuan’ and structure variants identified between yechensuan and Tunisia. A Features of the yechensuan genome. (i) coverage of illumine reads. (ii) coverage of Nanopore reads. (iii) GC content. (iv) Repeat content. (v) gene density (number of genes per Mb). B Structure variations between the yechensuan and Tunisia pomegranate genome. C Types and percentages of different transpositions detected in the yechensuan genome. D The percentage of position of structural variants relative to genes was showed in pie chart. E Size distribution of different types of structural variation. F Go term enrichment analysis of protein-coding genes affected by structure variation.

Alignment of the genome sequences of yechensuan and Tunisia showed good collinearity between the two reference genomes (Fig. 1B). Despite the high collinearity, 13 inversions and 11 translocations ranging from 760 bp to 1.69 Mb were identified. In addition to inversion and translations, indels larger than 50bp between yechensuan and Tunisia genomes were detected by direct comparison of the two assemblies. A total of 1,771 indels, ranging from 50 bp to 73.12 kb, were identified. The majority of the indels were relatively short with 71.48% smaller than 1 kb and only 4.6% larger than 10kb (Fig. 1E). Indels in gene body and upstream or downstream regions can impact gene functions and expression. Only 10% of identified indels overlapped with gene body or upstream or downstream (defined here as ±2 kb of gene body) regions (Fig. 1D). Gene ontology (GO) enrichment analysis showed that genes affected by the identified indels were enriched with those molecular function involved in ligase activity, enzyme activator, and peptide receptor activity (Fig. 1F).

In order to build a gene family-based pan-genome, we re-annotated the protein-coding genes of five other published *Punica granatum* genomes (Tunisia, Taishanhong, Dabenzi, Bhagwa, Azerbaijan) using the same annotation method. The number of protein-coding genes annotated in these 5 genomes ranged from 28,141 to 30,987, with an average of 29,133. The BUSCO evaluation for the protein-coding genes showed that 95-98% of the 1614 single copy Embryophyta genes were completely assembled in these genomes (Table 2), suggesting high completeness of the gene annotation. Orthofinder was used to identify orthologous gene families representing the genic content of the six pomegranate genomes. In total, we identified 28,207 gene families across the six genomes. Of those, 20,041 (71.05%) were present in all six genomes and were defined as core gene families. 7,922 (28.09%) were present in 2-5 genomes were defined as dispensable gene families. And the remaining 244 (0.86%) gene families were only present in one genome, defined as private genes (Fig 2B). All six accessions had accessions-specific orthogroups (Fig 2C). We found that the nucleotide diversity and Ka/Ks were higher in dispensable genes than in core genes (Fig 2E,F). 84.17 % core genes contained Pfams domains, which was much higher than the percentages in the dispensable and private genes (44.49% and 57.24%, respectively) (Fig 2D). These results indicated that core genes were more functionally conserved than dispensable genes. The number of gene families increased when including more genomes and approached a plateau when n = 6 (Fig 2A), suggesting that the six genomes are diverse and that a single reference genome cannot capture the full genetic diversity in pomegranate.

**Table 2.**
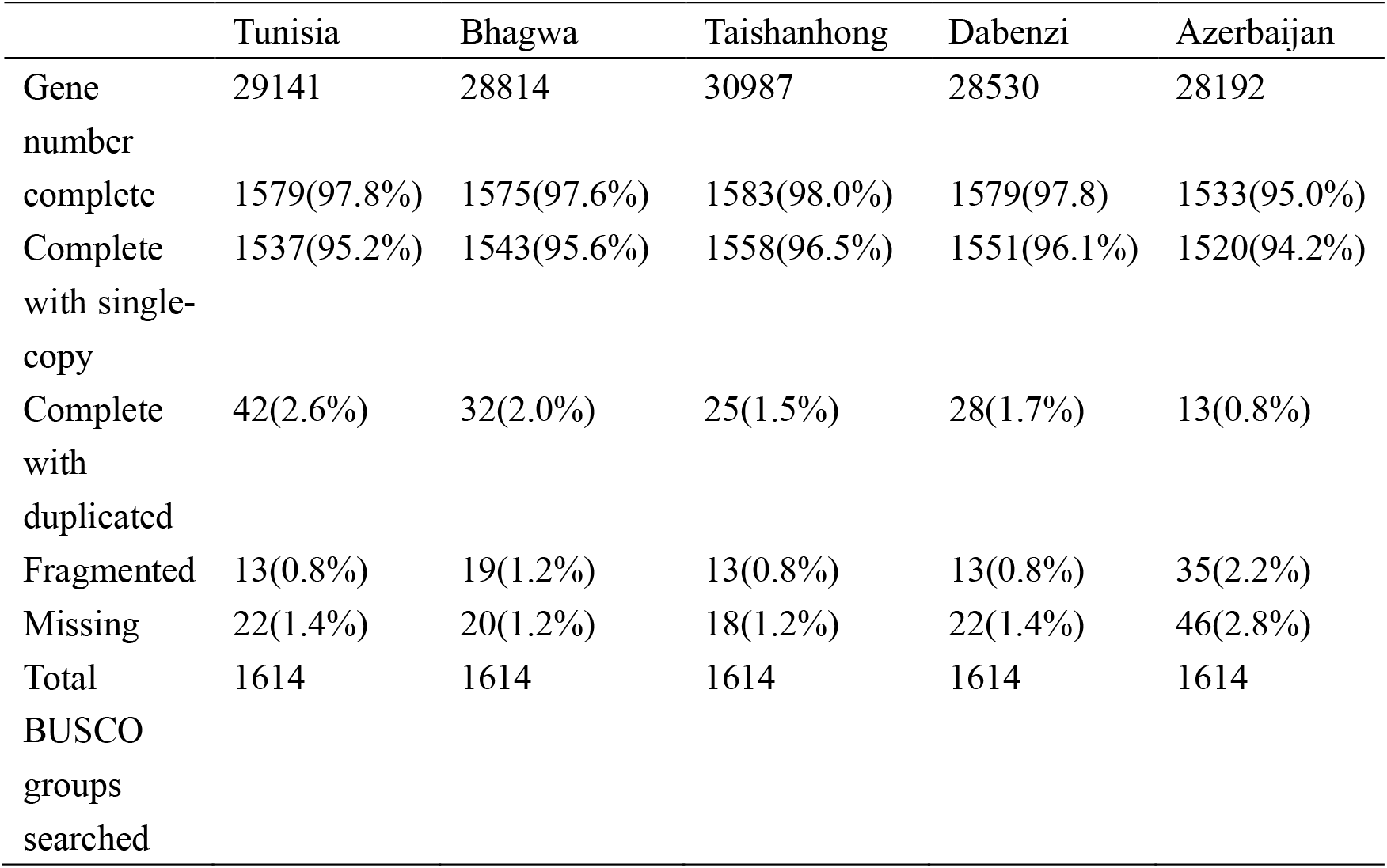
BUSCO evaluation summary of five pomegranate accession.

**Figure 2.**
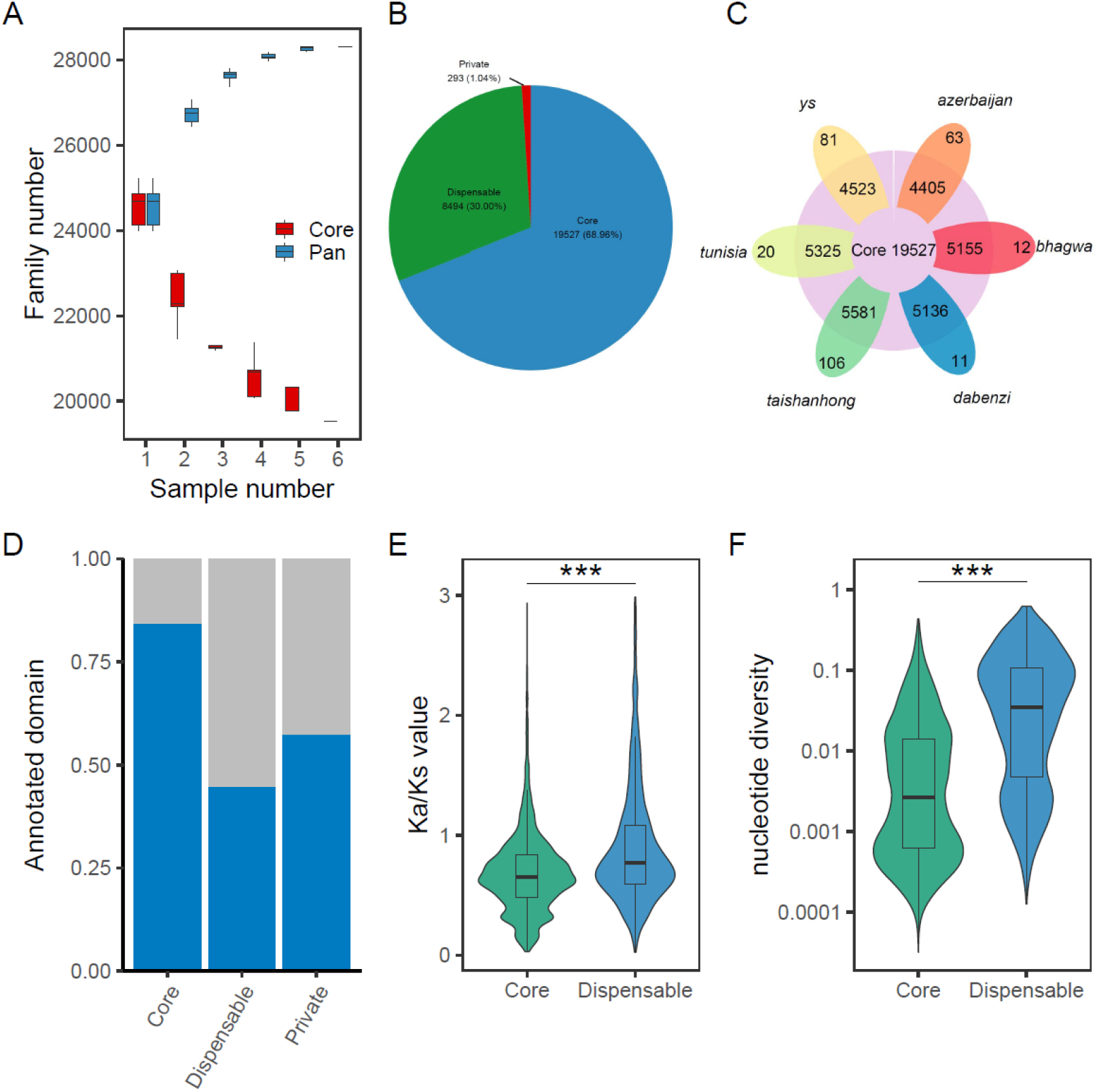
Pan- and Core Genome Analyses of 6 pomegranate accessions. A Variation of gene families in the pan-genome and core genome along with an additional pomegranate genome. B The Proportion of the gene family. C The flower plot displays the core orthogroups number (in the center), the orthogroups in a subset of accessions (in the annulus), and the accessions-specific orthogroups (in the petals) for the six pomegranate accessions. D Proportion of genes with Pfams domains in core, dispensable, and private genomes. Blue histograms indicate the genes with Pfams domains; gray histograms indicate the gene without Pfams domain. E and G Compositions of Ka/Ks value and nucleotide diversity in core and dispensable genes.

The genome assembly for sour pomegranate landraces provides a valuable set of resources for identifying causal variants that underlie traits for the domestication and improvement of pomegranate, serving as a basis for future research and crop improvement efforts for this important fruit crop.

## Methods

### Plant material

Punica granatum cv. ‘yechengsuan’ one-year-old branches were collected in its natural habitat from a single tree in Yecheng, Xinjiang, China. The branches were preserved in the experimental garden of Nanjing Forestry University by means of cuttings. After the cuttings grew leaves, we collected fresh leaves and frozen them immediately in liquid nitrogen for isolation of genomic DNA.

### DNA sequencing

For the Oxford Nanopore library preparation, a total of 10 ug high-molecular-weight DNA was extracted for library preparation according to the manufacturer’s instructions. The resulted library was combined with RBF (ONT, Oxford, UK) and loaded onto primed R10.3 Flow cells (Oxford Nanopore Technologies) for sequencing on a PromethION high-throughput platform according to the manufacturer’s recommendations. Base calling analysis was performed in real-time on the compute module (Guppy4.0.11). only passed reads with a Q-value threshold of seven were used for further data analysis. We also generate ∼40X Illumina paired-end reads on the HiSeq 2000 platform (Illumina, San Diego, CA, USA) with insert sizes of 350 bp for polishing the genome assembly. Raw reads were processed using fastp ^18^ for filtering poor quality reads with default parameters.

### Estimation of the genome size and heterozygosity

The genome size of Punica granatum cv ‘yechengsuan’ was estimated using GenomeScope ^19^. The quality of the Illumina reads was estimated using the FastQC program. The analysis of optimal kmer size was performed by using KmerGenie ^20^ with chloroplast and mitochondrial sequences being removed from the high-quality clean reads. Then, the best k-mer was used for kmer count analysis with Jellyfish ^21^. After converting the k-mer counts into a histogram format, the k-mer distribution was analyzed to estimate genome size and heterozygosity.

### Genome assembly

The long-read assembler NextDenovo ^22^ was used to assemble the ONT long reads with parameters: ‘read_cutoff = 1k’ and ‘seed_cutoff = 49,111’. NextPolish ^23^ was used to polish the contigs assembled by ONT long reads with illumina paired-end reads. Pseudochromosomes were constructed according to a reference-based approach using Ragtag ^24^ with default parameters based on the ‘Tunisia’ reference genome. TGS-gapcloser ^25^ was used to fill gaps.

### Repetitive element annotation

The library of species-specific repeats was constructed using RepeatModeler ^26^ with default options. DeepTE ^27^ was then used to reclassify the TE library that were annotated as “LTR/unknown” by RepeatModeler. RepeatMasker ^28^ was used to identify repeat elements with the specific library and the default library from the RepeatMasker database.

### Prediction and annotation of protein-coding genes

Protein-coding genes were annotated by integrating ab initio prediction, homology searches, and mRNA expression evidence. Two ab initio gene prediction tools were used including Augustus ^29^, and GeneMark ^30^. Predicted proteins from four plant genomes (*Arabidopsis thaliana, Malus domestica, Vitis vinifera*, and *Eucalyptus grandis*) were used to perform homology-based prediction with miniport ^31^. As for the third approach, RNA-seq data from different tissues were download from NCBI SRA database and then were assembled using HISAT2 ^32^ and stringtie ^33^. Transdecoder was then used to infer the structures of gene models. By giving weights for the three methods, all predicted gene structures were synthesized into consensus gene models using EVidenceModeler ^34^. We re-annotated the protein-coding genes of five other published pomegranate genomes using the same annotation method. For gene function annotation, eggnog-mapper ^35^ was applied to get seed ortholog and function description, Gene Ontology (GO) number, Enzyme Commission nomenclature (EC) number, Kyoto Encyclopedia of Genes and Genomes (KEGG) number, PFAM number and so on.

### Pan-genome construction

Ortholog groups among the six pomegranate accession genomes were identified using OrthoFinder ^36^ with default parameters. The resulting gene families were divided into core, dispensable, and private based on the number of genomes contained in each cluster. Gene families present in all of the genomes were defined as core. Gene families present in 2-5 genomes were defined as dispensable. Gene families present only in one genome were defined as private. Single-copy ortholog groups containing more than two accessions were used to calculate ka/ks and nucleotide diversity. Protein sequences were first alignment by MAFFT ^37^ and then converted to DNA codon alignments using ParaAT ^38^. Kaks_calculator 3.0 ^39^ was used to estimate the Ka/Ks value for each ortholog group. Nucleotide diversity was calculated with R function nuc.div in pegas package ^40^.

## Data record

All sequence data generated for this study were deposited in NCBI, under the BioProject PRJNA982616. The genome assembly in this study are available at Genbank database under the BioProject PRJNA1034978. All six pomegranate accession genomes and corresponding genome annotation files are available at (https://doi.org/10.6084/m9.figshare.24476632.v1)

### Technical validation

The completeness of the assembled genome sequence was estimated by BUSCO ^41^ using the embryophyte_odb10 database with default parameters. The BUSCO analysis showed that 98.8% of the expected embryophyte_odb10 genes (single-copy gene: 96.6% and duplicated genes: 2.2%) were identified as complete, and 0.2% fragmented genes were found in the genome assembly. The LAI (LTR Assembly Index) calculated with LTR_retriever ^42^ was used to assess the assembly completeness of intergenic regions. The LAI value was 12.30, which indicated that out newly assembly reached the ‘reference’ level (LAI > 10). The evaluations indicated high continuity and high completeness of our newly assembly. The completeness of the gene annotation was also evaluated using BUSCO with completeness scores 97.4%, suggesting high gene annotation quality. Around 90% of the genes were functionally annotated through at least one of the databases in eggnog.

### Code availability

The code used in this study are available through github (https://github.com/NotebookOFXiaoMing/sourpomegenome)

## Acknowledgements

The authors acknowledge the Research/Scientific Computing teams at The James Hutton Institute and NIAB for providing computational resources and technical support for the “UK’s Crop Diversity Bioinformatics HPC” (BBSRC grant BB/S019669/1), use of which has contributed to the results reported within this paper. This work was supported by the Initiative Project for Talents of Nanjing Forestry University (GXL2014070, GXL2018032), the Priority Academic Program Development of Jiangsu High Education Institutions (PAPD), the Natural Science Foundation of Jiangsu Province (BK20180768), the National Nature Science Foundation of China (31901341), China Scholarship Council (CSC, 202108320293), Postgraduate Research & Practice Innovation Program of Jiangsu Province (KYCX20_0874)

## Author contributions

M.Y. assisted with data analysis and wrote the manuscript. X.Q.Z., and L.Y. processed data and assisted with data analysis. Z.H.Y. supervised all aspects of the study and wrote the manuscript. All authors participated in careful editing of manuscript.

